# Alterations in Acylcarnitines, Amines, and Lipids Inform about Mechanism of Action of Citalopram/Escitalopram in Major Depression

**DOI:** 10.1101/2020.02.10.927012

**Authors:** Siamak MahmoudianDehkordi, Ahmed T. Ahmed, Sudeepa Bhattacharyya, Xianlin Han, Rebecca A. Baillie, Matthias Arnold, Michelle K. Skime, Lisa St. John-Williams, M. Arthur Moseley, J. Will Thompson, Gregory Louie, Patricio Riva-Posse, W. Edward Craighead, William McDonald, Ranga Krishnan, A John Rush, Mark A. Frye, Boadie W. Dunlop, Richard M. Weinshilboum, The Mood Disorders Precision Medicine Consortium (MDPMC), Rima Kaddurah-Daouk

**Author notes:** Corresponding author: Rima Kaddurah-Daouk, Department of Psychiatry, Duke University, Durham, NC, USA. co-first authors (Siamak MahmoudianDehkordi, Ahmed T. Ahmed). **Abbreviations:** Acylcarnitines abbreviations for names of all metabolites measured using the P180 platform can be found in Supplementary Table 1.

## Abstract

Selective serotonin reuptake inhibitors (SSRIs) are the first-line treatment for major depressive disorder (MDD), yet their mechanisms of action are not fully understood and their therapeutic benefit varies among individuals. We used a targeted metabolomics approach utilizing a panel of 180 metabolites to gain insights into mechanisms of action and response to citalopram/escitalopram. Plasma samples from 136 participants with MDD enrolled into the Mayo Pharmacogenomics Research Network Antidepressant Medication Pharmacogenomic Study (PGRN-AMPS) were profiled at baseline and after 8 weeks of treatment. After treatment, we saw increased levels of short-chain acylcarnitines and decreased levels of medium- and long-chain acylcarnitines, suggesting an SSRI effect on β-oxidation and mitochondrial function. Amines – including arginine, proline, and methionine-sulfoxide – were upregulated while serotonin and sarcosine were downregulated, suggesting an SSRI effect on urea cycle, one-carbon metabolism and serotonin uptake. Eighteen lipids within the phosphatidylcholine (PC aa and ae) classes were upregulated. Changes in several lipid and amine levels correlated with changes in 17-item Hamilton Rating Scale for Depression scores (HRSD_17_). Differences in metabolic profiles at baseline and post-treatment were noted between participants who remitted (HRSD_17_≤7) and those who gained no meaningful benefits (<30% reduction in HRSD_17_). Remitters exhibited a) higher baseline levels of C3, C5, alpha-aminoadipic acid, sarcosine and serotonin; and b) higher week-8 levels of PC aa C34:1, PC aa C34:2, PC aa C36:2 and PC aa C36:4. These findings suggest that mitochondrial energetics – including acylcarnitine metabolism, transport and its link to β-oxidation – and lipid membrane remodeling may play roles in SSRI treatment response.

## Introduction

Major depressive disorder (MDD) is a common, often disabling condition that affects more than 300 million individuals worldwide ^1^, but much about its pathobiology and the biology of treatment response remains unknown. Selective serotonin reuptake inhibitors (SSRIs) are common first-line agents used for the treatment of MDD ^2, 3^, yet roughly 40% of patients who receive SSRI treatment do not respond and more than two thirds do not achieve remission of symptoms ^4^. Clinical symptoms are insufficient to guide treatment selection for individual patients ^5^. Hence, in clinical practice a “trial and error” approach is used to find an effective therapy ^6^.

MDD patients who achieve remission after treatment with an antidepressant appear to have a metabolic state distinct from never-depressed individuals, with alterations in methylation, purine metabolism and oxidative stress pathways ^7^. Several pilot studies have implicated metabolic dysregulation in the pathogenesis of MDD, such as alterations in several pathways, including neurotransmission (GABA, glutamine, tryptophan, phenylalanine), nitrogen metabolism, methylation, and lipid metabolism ^8–12^.

A metabolomics approach provides tools to enable the mapping of global metabolic changes in neuropsychiatric diseases and upon treatment ^7–9, 13–18^. Pharmacometabolomics – the application of metabolomics to the study of drug effects – has been successfully used to map effects of sertraline ^19^, ketamine, and placebo ^20^ by providing new insights about mechanisms of action and response ^19, 21, 22^.

Several studies have shown mitochondrial dysfunction or lower adenosine triphosphate (ATP) production in MDD patients, which suggests a role for mitochondrial energetics in depression ^23–27^. Furthermore, accumulated evidence suggests that alterations of acylcarnitines may contribute to neuropsychiatric diseases including depression ^28–32^, autism ^33^ and schizophrenia ^34, 35^. Acylcarnitines are a class of metabolites that are formed from the transfer of the acyl group of a fatty acyl-Coenzyme A (CoA) to carnitine, which is catalyzed by carnitine acyltransferases in mitochondria (**Fig. 1A**) ^36–38^. Carnitine deficiency or dysfunction in the carnitine acyltransferase results in a reduced β-oxidation of fatty acids, and therefore reduced mitochondrial energy (ATP) production. Hence, altered plasma or serum acylcarnitine levels can be used as biomarkers that suggest abnormalities in beta-oxidation (e.g., inborn errors of metabolism) ^39–43^. Elevated medium- and long-chain acylcarnitine concentrations in blood have been associated with incomplete β-oxidation of fatty acids in a rat model of depression ^44^.

**Fig.1:**
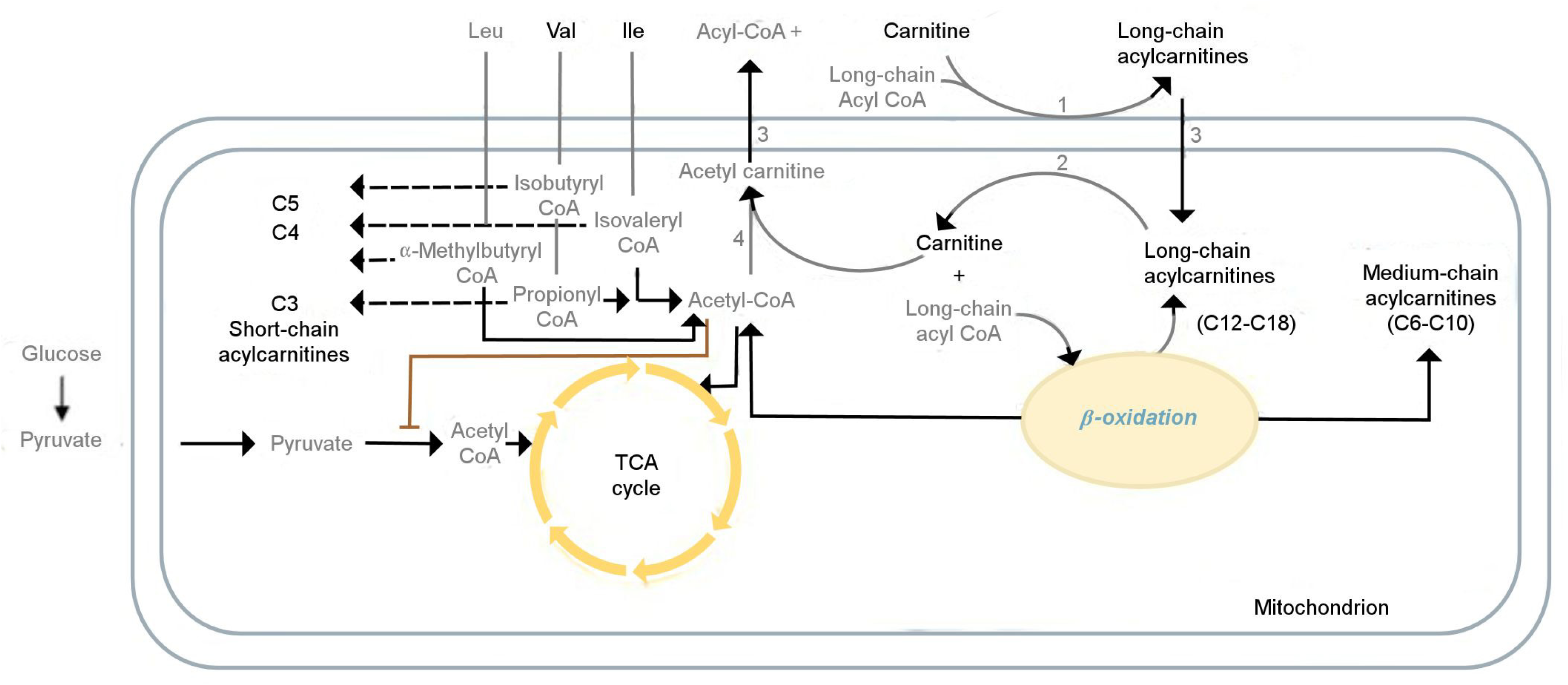
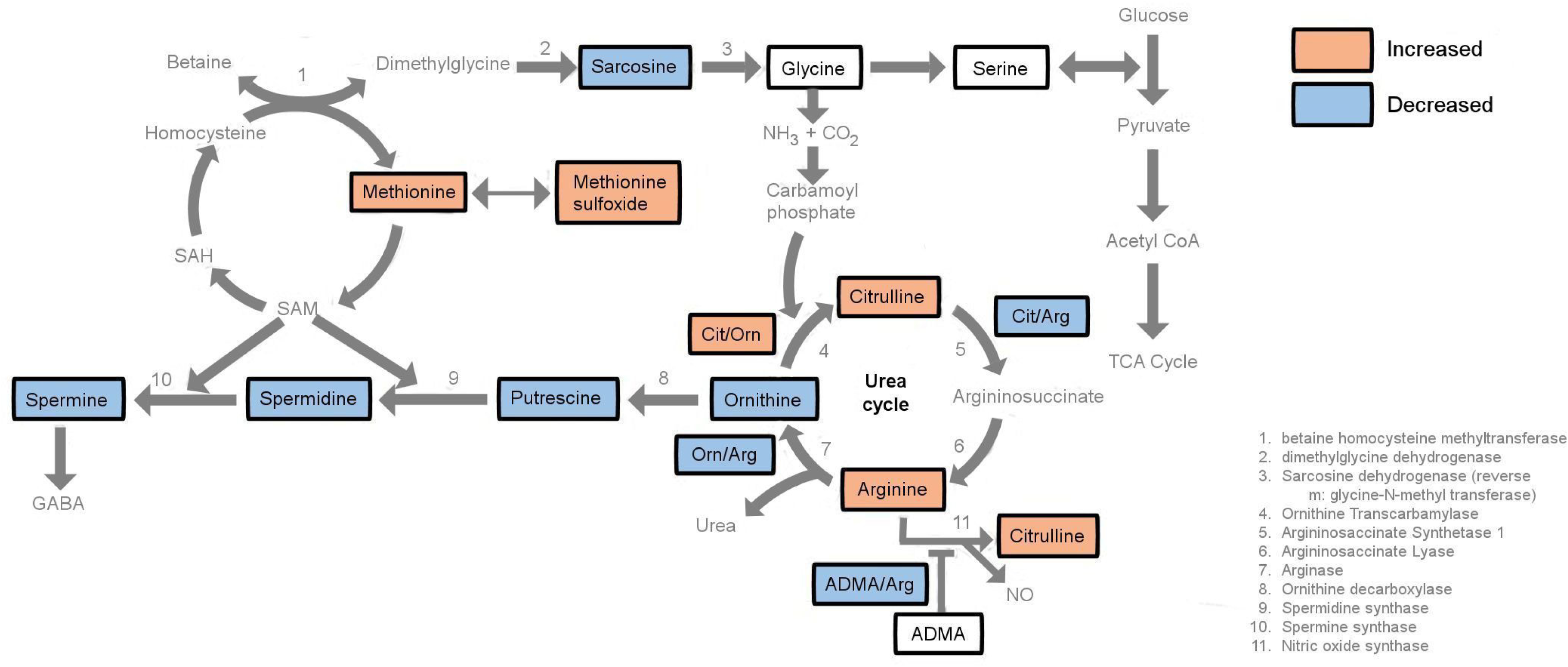
Metabolism Pathways and the Effect of SSRIs. **Fig. 1A: Role of acylcarnitines in β-oxidation of fatty acids** Mitochondrial energy metabolism pathways involve: branched-chain amino acids (BCAAs), acyl carnitines, fatty acid beta-oxidation and tricarboxylic acid cycle (TCA cycle). A delicate balance between two of the main cellular energy resources, glucose utilization and fatty acid utilization, is essential to maintain physiological homeostasis, the perturbation of which may lead to pathologic processes. The grey font metabolites have not been measured in p180. *Abbreviations*: Leu: Leucine, Val: Valine, Ile: Isoleucine, CoA: Coenzyme A, CPT1: carnitine-palmitoyl-transferase 1, CPT2: carnitine-palmitoyl-transferase 2, CACT: carnitine-acylcarnitine-translocase, CrAT: carnitine O-acetyltransferase. Metabolite abbreviations are spelled out in Supplementary Table 1. **Fig. 1B: Effect of SSRI on interconnected pathways, including polyamine, sarcosine, urea cycle and 1-carbon metabolism** Orange boxes indicate an increase in metabolite levels, while blue boxes represent a decrease in metabolite levels. White boxes represent no change in in metabolite levels. Metabolites not measured in this assay are in grey. *Abbreviations*: NH_3_: Ammonia, CO_2_: Carbon Dioxide, SAH: S-adenosyl-l-homocysteine, SAM: S-adenosyl methionine, Cit: Citrulline, Orn: Ornithine, Arg: Arginine, ADMA: Asymmetric dimethylarginine, CoA: Coenzyme A.

Changes in the levels of amino acids and biogenic amines such as histidine, kynurenine, methionine sulfoxide, arginine, citrulline, ornithine and urea have been implicated in MDD ^45–53^. Moreover, lipids are involved in crucial brain functions including cell membrane structure, membrane transmitters and regulation of synapses, as well as biological messenger functions, energy metabolism and neuroendocrine function ^54^. Blood lipid profile changes have been implicated in the pathogenesis of depression, schizophrenia, and Alzheimer’s disease ^55–59^. MDD entails disturbances in the regulation of the molecular pathways such neurotransmitter systems, synaptic plasticity, and the neuroendocrinological and immune regulator ^60^. Therefore, lipid analysis may play a critical role in finding MDD-relevant biomarkers ^61^.

In this study, we used a targeted metabolomics approach to interrogate possible functions for acylcarnitines, amino acids, biogenic amines, and lipids to addresses the following questions regarding mitochondrial function and neurotransmission in MDD:

1. What are the overall changes in the metabolic profile over 8 weeks of SSRI exposure, and which of these metabolic changes are related to each other?
2. Which metabolite level changes are related to changes in depressive symptoms (17-item
3. Rating Scale for Depression [HRSD_17_]) over the 8 weeks of SSRI treatment?
4. Are there baseline metabolites that differentiate between patients who have benefitted substantially (HRSD_17_≤7) after 8 weeks of treatment vs. those who gained no meaningful benefits with SSRI treatment (<30% reduction in HRSD_17_ from baseline to week 8)?
5. Are there week 8 metabolites that differentiate between those who have benefitted substantially vs. those who gained no meaningful benefits?

## Materials and Methods

### Study Design and Participants

This study examined plasma samples from 136 participants with MDD who were enrolled into the Mayo Pharmacogenetics Research Network Antidepressant Medication Pharmacogenetics Study (PGRN-AMPS) (ClinicalTrials.gov NCT00613470). The design and clinical outcomes of the PGRN-AMPS study have been previously published ^62^. The trial enrolled 800 MDD participants 18-84 years of age from Mayo Clinic psychiatry or primary care clinics. Participants received 8 weeks of open-label treatment with either citalopram (20-40 mg/day) or escitalopram (10-20 mg/day). MDD severity was assessed using the HRSD_17_ ^63^ at each visit. The PGRN-AMPS protocol was approved by the Mayo Clinic Institutional Review Board. All risks and benefits of the PGRN-AMPS study were discussed with participants, each of whom gave written informed consent prior to entering the study.

### Metabolomic Profiling using the Absolute IDQ p180 Kit

Using the AbsoluteIDQ® p180 Kit (BIOCRATES Life Science AG, Innsbruck, Austria), we measured metabolites with a targeted metabolomics approach using an ultra-performance liquid chromatography (UPLC)/MS/MS system (Acquity UPLC [Waters], TQ-S triple quadrupole MS/MS [Waters]). This system provides measurements of up to 180 endogenous metabolites from various classes including acylcarnitines, amino acids, biogenic amines, glycerophospholipids and sphingolipids. The AbsoluteIDQ® p180 kit has been validated according to European Medicine Agency Guidelines on bioanalytical method validation. Further, the kit plates include an automated technical validation to assure the validity of the run and provide verification of the actual performance of the applied quantitative procedure, including instrumental analysis. Using MetIDQ® software, we performed the technical validation of each analyzed kit plate, which is based on results obtained and defined acceptance criteria for blank, zero samples, calibration standards and curves, low/medium/high-level quality control (QC) samples, and measured signal intensity of internal standards over the plate. Several previous studies have used the same platform for MDD participants ^20, 52, 64^. We analyzed de-identified samples following the manufacturer’s protocol, with metabolomics labs blinded to clinical data.

### Quality Control of P180 Profiles

The raw metabolomic profiles included 182 metabolite measurements from 578 plasma samples. Each assay plate contained a group of duplicates that were acquired by combining roughly 10 μl from the first 76 samples in the study (QC pool duplicates) to allow for appropriate inter-plate abundance scaling based specifically on this cohort of samples (n=24 across all plates). Metabolites with >40% of measurements below the lower limit of detection (LOD) were excluded from the analysis (n=163 metabolites remained in the analysis). Imputation of <LOD values was performed using each metabolite’s LOD/2 value. To adjust for the batch effects, a correction factor for each metabolite in a specific plate was calculated by dividing metabolites’ QC global average by QC average within the plate. Metabolite concentrations were log2 transformed for statistical analysis. Individuals having both a baseline and a week 8 sample were kept for the subsequent statistical analysis (n=136 participants). This resulted in an analysis data set containing 377 samples (n=136 at baseline, n=105 at week 4, and n=136 at week 8) and 163 metabolites.

### Clinical Outcomes

The HRSD_17_ total score was used as the outcome measure for both continuous depression symptom severity change and for defining categorical outcomes. Consistent with prior definitions ^65, 66^, participant outcomes at week 8 were categorized as “remitters” (HRSD_17_≤7), “response without remission” (≥50% reduction from baseline HRSD_17_, but not reaching remission threshold); “partial response” (30-49% reduction from baseline HRSD_17_ score); and “treatment failures” (<30% reduction from baseline HRSD_17_ score).

### Statistical Analysis

Differences in demographic variables and depression scores across the response groups were evaluated using ANOVA and the Pearson Chi-squared test (for categorical variables). All association and differential abundance analyses were performed in a metabolite-wise manner. To examine the significance of log2-fold change in metabolite concentrations, linear mixed effect models (with random intercept) with log2 metabolite levels as the dependent variable were fitted while correcting for age, sex, baseline HRSD_17_ and specific antidepressant (citalopram/escitalopram). Then we used the “emmeans” R package to compute the least squared means of the contrasts of interest (week 8 vs. baseline) and their corresponding *p*-values. Adjustments for multiple comparisons were made using the Benjamini-Hochberg procedure to control the false discovery rate. We investigated the global correlation structure of changing metabolites from baseline to week 8 using Spearman’s ranked correlation to identify biochemically related metabolites on which SSRI exposure has similar effect, followed by hierarchical clustering to group similar correlated metabolites.

To detect whether changes in metabolites were associated with clinical outcome, we conducted continuous and categorical analyses. In the continuous analysis, the associations of changes in HRSD_17_ score after 8 weeks with changes in metabolite levels were tested using linear regression models corrected for age and sex. In the categorical analysis, profiles of remission (“remitters” vs. “response” vs. “partial response” vs. “treatment failure”) were compared using linear mixed effect models (with random intercept), with log2 metabolite levels as the dependent variable and the interaction of the week 8 outcome (4 level categorical variable: remitters; response without remission; partial response; treatment failures) and visit (2 level categorical variable: baseline; week 8) as independent variables while correcting for age, sex, HRSD_17_ score and antidepressant (citalopram/escitalopram), as the method used by other investigators to maximize the ability to identify biological characteristics most clearly associated with differential outcomes ^66 67^. Then the fitted models were used to conduct contrasts between the “remission” and “treatment failure” groups at baseline and week 8 using the “emmeans” R-package.

## Results

Demographic and clinical features of the 136 participants with MDD whose plasma was analyzed in the present study are summarized in **Table 1**. At baseline, age and sex were not significantly different across response groups. Metabolites measured in the p180 kit, as well as QC summary statistics, can be found in **Supplementary Table 1**.

**Table 1.** Clinical and Demographic Characteristics of Mayo PGRN-AMPS Study Participants Stratified by Remission Status

| Characteristic | N | Grand Mean | Remission<sup>a</sup> (N=64) | Response without Remission (N=30) | Partial-Response (N=30) | Treatment Failure (N=12) | Test Statistic |
| --- | --- | --- | --- | --- | --- | --- | --- |
| Age (years), Mean (SE) | 136 | 40.17(13.60) | 40.76(1.72) | 35.58 (1.91) | 41.31(2.26) | 45.75(5.63) | F=1.4<br>d.f.=3,132<br>P=0.24 |
| Sex: Male | 136 | 36% (49) | 34% (22) | 30% ( 9) | 29% ( 13) | 50% ( 6) | Chi-square=1.8<br>d.f.=3<br>P=0.62 |
| Body Mass Index (kg/m<sup>2</sup>) | 97 | 28.77(6.52) | 29.19(6.58) | 26.58(5.43) | 29.42(7.12) | 29.54(6.83) | F=1<br>d.f.=3,93<br>P=0.39 |
| HRSD<sub>17</sub> at Baseline Mean (SD) | 136 | 23.38(4.91) | 22.76(0.66) | 25.70(0.85) | 22.83(0.70) | 22.25(1.30) | F=214<br>d.f.=3,132<br>**P<0.001** |
| HRSD<sub>17</sub> at Week 8 Mean (SD) | 136 | 8.74(5.49) | 4.06(0.27) | 10.10(0.37) | 13.43 (0.47) | 18.50(1.15) | F=3.4<br>d.f.=3,132<br>**P=0.018** |

Remission status based on week-8 HRSD<sub>17</sub> score. BMI measurement is available only at baseline.
*Abbreviations:* PGRN-AMPS: Mayo Pharmacogenomics Research Network Antidepressant Medication Pharmacogenomic Study, BMI: Body Mass Index, HRSD<sub>17</sub>: The 17-item Hamilton Rating Scale for Depression.

### What are the Overall Changes in the Metabolic Profile over 8 Weeks of SSRI Exposure, and Which of these Metabolic Changes are Related to Each Other?

After 8 weeks of SSRI treatment, levels of several metabolites from different classes were changed significantly (*q*-value <0.05; see **Fig. 2A** and **Table 2**). Among the acylcarnitines, we observed an increased level of 3 short-chain acylcarnitines (C3, C4 and C5) and a decrease in medium and long-chain acylcarnitines (e.g., C8, C10, C12, C14:2, C16, C16:1, C18, C18:1, and C18:2). Within the classes of amino acids and biogenic amines, we observed a statistically significant increase in two amino acids (arginine and proline) and one biogenic amine (methionine sulfoxide), and a significant decrease in two biogenic amines (serotonin and sarcosine). Among the lipid classes, we observed a statistically significant up regulation of several phosphatidylcholines (PCs) (e.g., PC aa-C36:1,-C30:0,-C42:2; ether phosphatidylcholines, PC ae C34:3, -C38:2,-C36:3) and one sphingolipid (sphingomyelin [SM] C24:0).

**Fig 2:**
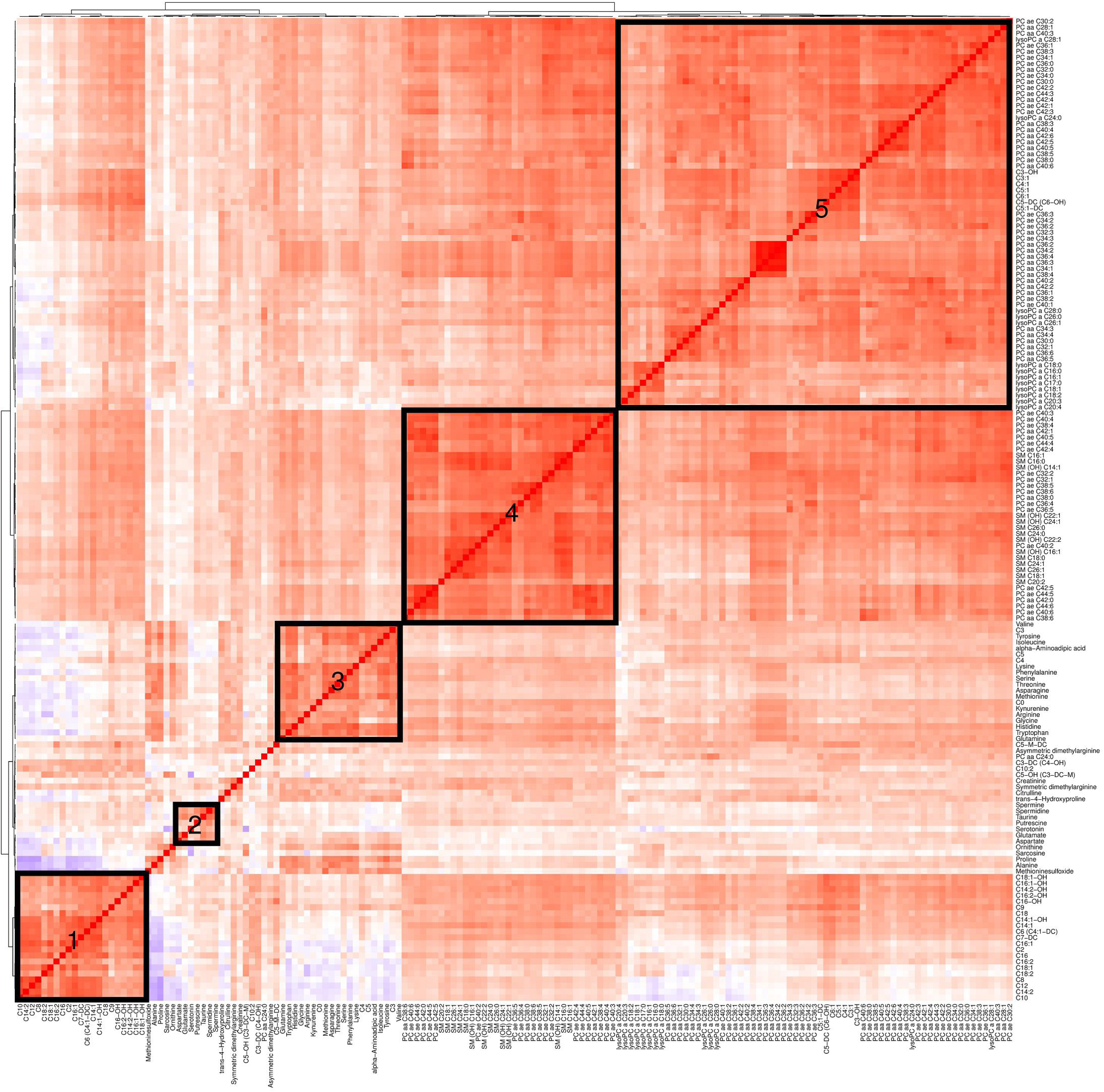
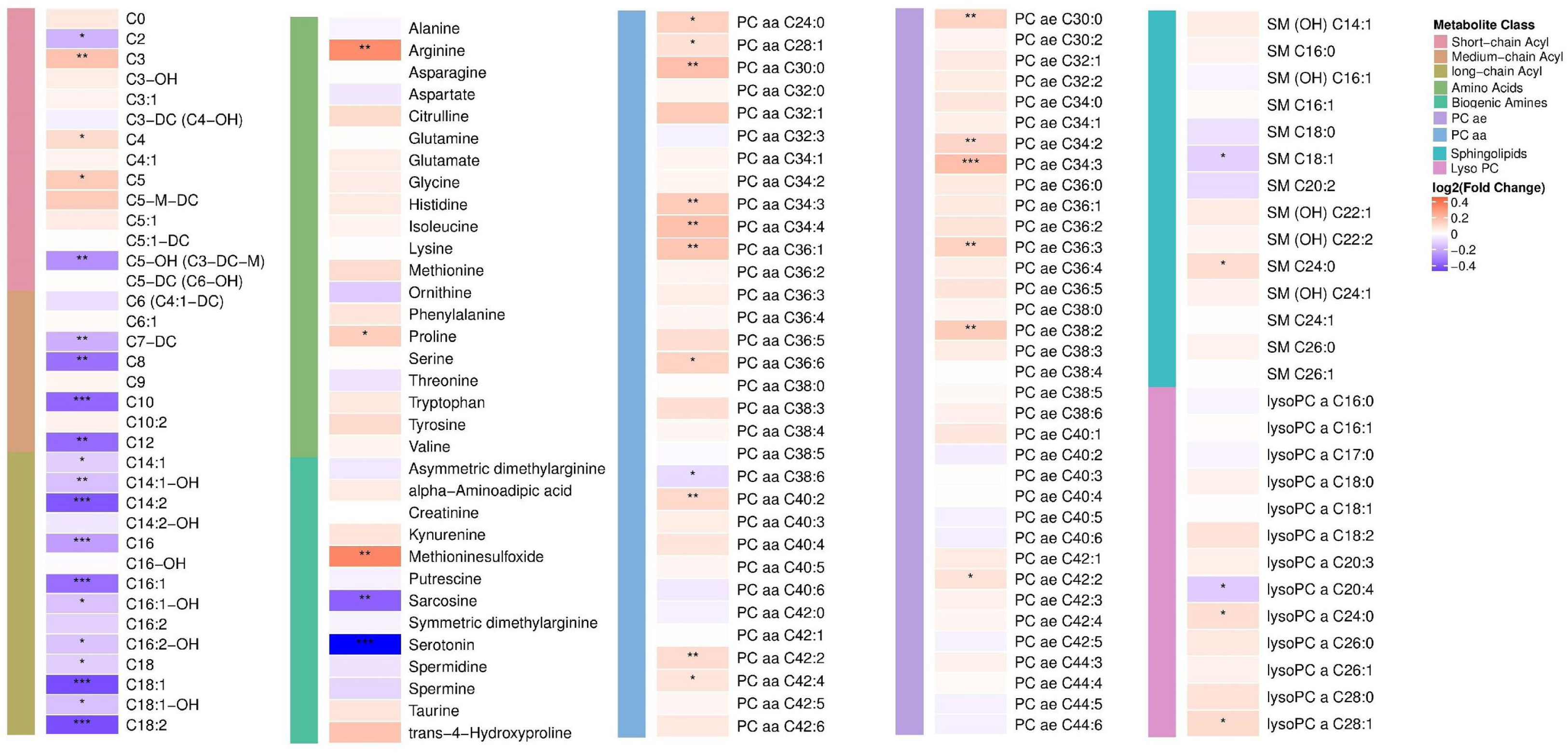
Change in Metabolite Levels from Pre-treatment to 8 Weeks Post-treatment with Escitalopram/Citalopram. **Fig. 2A: Heat map depicting log2-fold changes in metabolite levels** P-values were obtained using linear mixed effect models controlling for age, sex and baseline HRSD_17_ were corrected for multiple comparisons. Red indicates an increase and blue indicates a decrease in metabolite levels over 8 weeks of treatment; *: *q*-value<0.05, **: *q*-value <0.01 and ***: *q*-value <0.001. *Abbreviations*: PC: Phosphatidylcholine, SM: Sphingomyelin, LysoPC: Lyso-Phosphatidylcholines, SSRI: Selective Serotonin Reuptake Inhibitor, HRSD_17_: 17-item Hamilton Rating Scale for Depression. Metabolite abbreviations are spelled out in Supplementary Table 1. **Fig. 2B: Hierarchical clustering of Spearman’s rank correlation of change in metabolite levels** Red represents positive correlations and blue represents negative correlations. Cluster #1: enriched for medium and long-chain acylcarnitines (e.g., C8, C9, C10, C12, C16), Cluster #2: biogenic amines including serotonin, spermine, spermidine, taurine, putrescine, glutamate, ornithine and sarcosine. Cluster #3: short chain–acylcarnitines (C3, C4, C5), branched-chain amino acids (isoleucine and valine), kynurenine, arginine, glycine, tryptophan and glutamine. Cluster #4: long-chain phosphodelcholine of PC ae class and Cluster #5: a large number of phospholipids and lyso-phospholipids. *Abbreviations*: PC: Phosphatidylcholine, LysoPC: Lyso-Phosphatidylcholines, SM: Sphingomyelin. Metabolite abbreviations are spelled out in Supplementary Table 1.

**Table 2.**
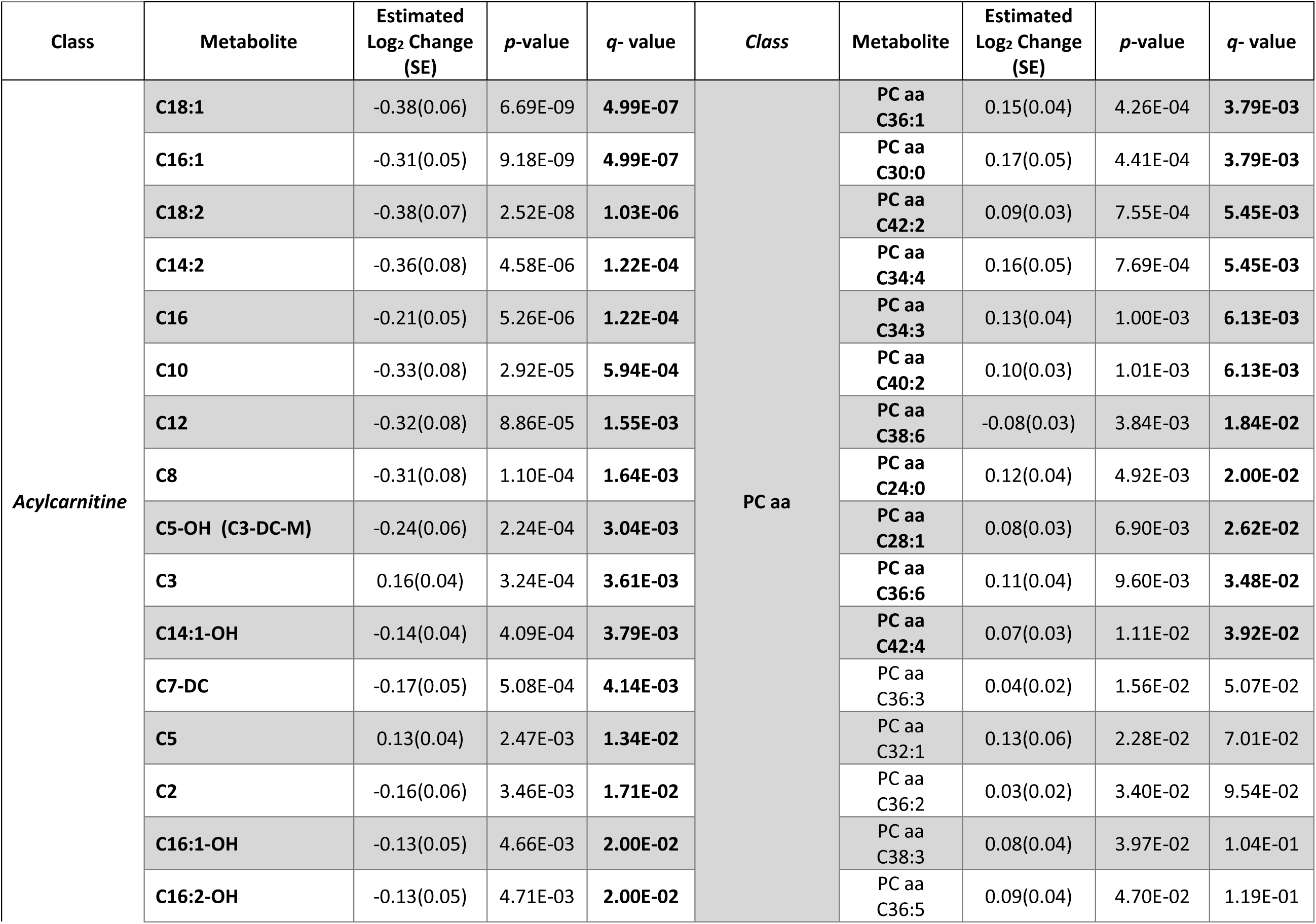
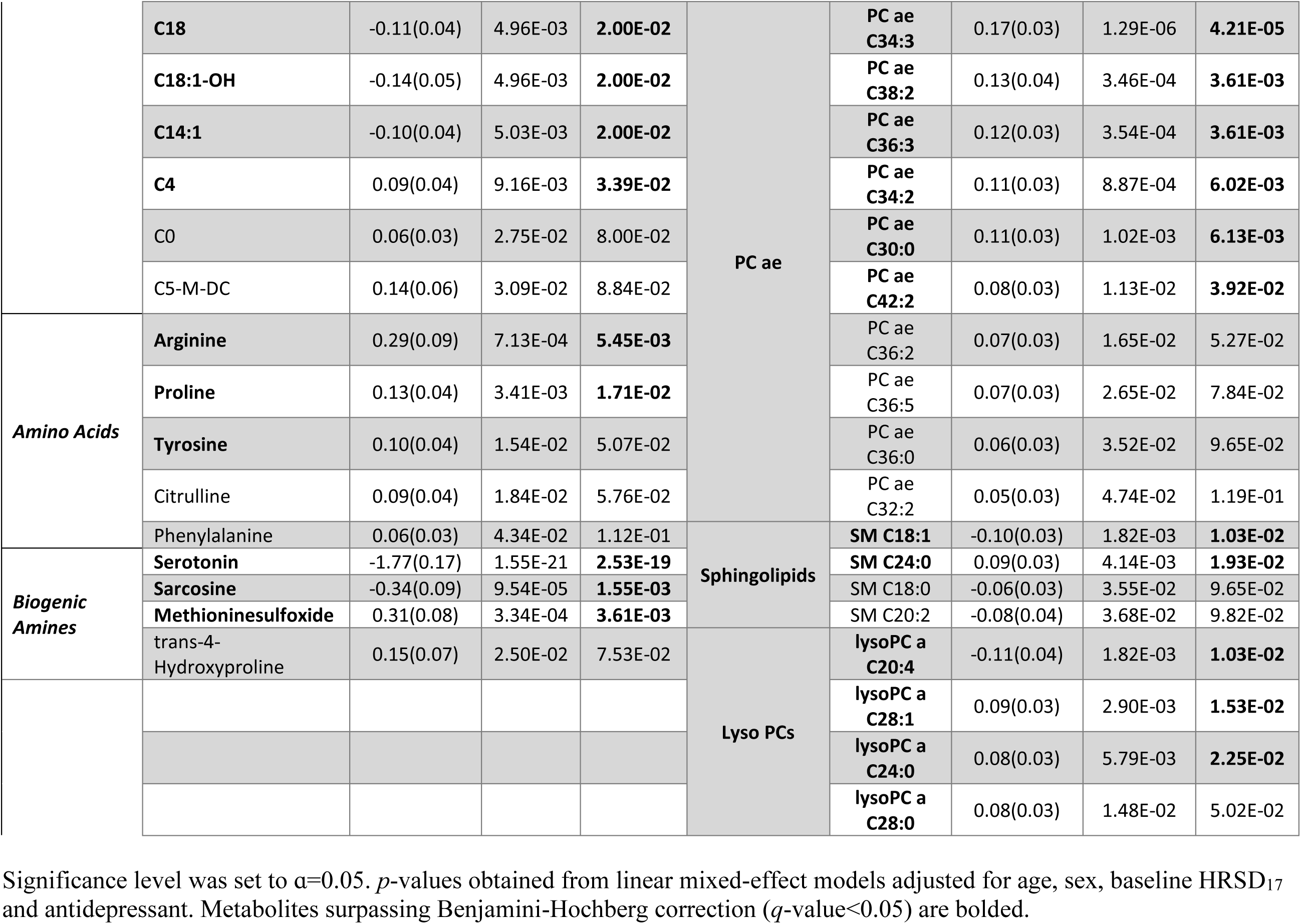
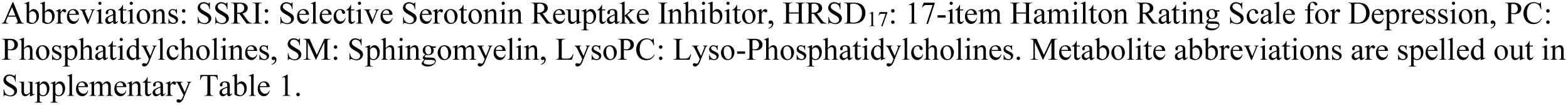
Statistically Significant changes in Metabolite Levels after 8 weeks of SSRI Treatment.

As depicted in **Fig. 2B**, correlation analysis revealed that several metabolite changes in response to the drug exposure were highly correlated, especially notable amongst metabolites belonging to the same class. We observed five distinct clusters: cluster 1) mainly formed with medium- and long-chain acylcarnitines (e.g., C8, C9, C10, C12, C16); cluster 2) biogenic amines including serotonin, spermine, spermidine, taurine, putrescine, glutamate, ornithine and sarcosine; cluster 3) short-chain acylcarnitines (C3, C4, C5), branched-chain amino acids (isoleucine and valine), kynurenine, arginine, glycine, tryptophan and glutamine; cluster 4) long-chain ether phospholipids; and cluster 5) a large number of phospholipids and lyso-phospholipids.

### Which Metabolite-level Changes are Related to Changes in Depressive Symptoms (HRSD_17_) over 8 Weeks of SSRI Treatment?

We examined the association of change in HRSD_17_ with log2-fold change in metabolite levels from baseline to week 8, adjusting for the covariates of age and sex. The associations with uncorrected *p*-value <0.05 included: one acylcarnitine (C5-M-DC), four amines (histidine, proline, kynurenine and trans5 hydroxyproline), seven PC aas and seven PC aes. Change in all of these metabolites were inversely associated with change in HRSD_17_ (**Supplementary Table 2)**.

### At baseline and After 8 Weeks of Treatment, Which Metabolomic Profiles Differentiate Between Remitters vs. Treatment Failures?

We compared the remitter (n=64) and treatment failure (n=12) groups at baseline and after 8 weeks of SSRI treatment. Eleven metabolites were significantly different (unadjusted *p*-value <0.05) between the two groups at either baseline or week 8 (**Fig. 3 and Supplementary Table 3**). Levels of two short-chain acylcarnitines (C3 and C5) and two biogenic amines (alpha-amino adipic acid and sarcosine) were higher at baseline in the remitters and remained higher at the end of treatment. Serotonin and C3 were significantly different at baseline, but not at week 8. SSRI treatment resulted in differential regulation of four phosphatidylcholines (PC aa C34:2, PC aa C36:2, PC aa C36:4 and PC aa C43:1) and two lysoPCs (lysoPC a C18:2 and lysoPC a C20:4) between the two groups at week 8.

**Fig. 3:**
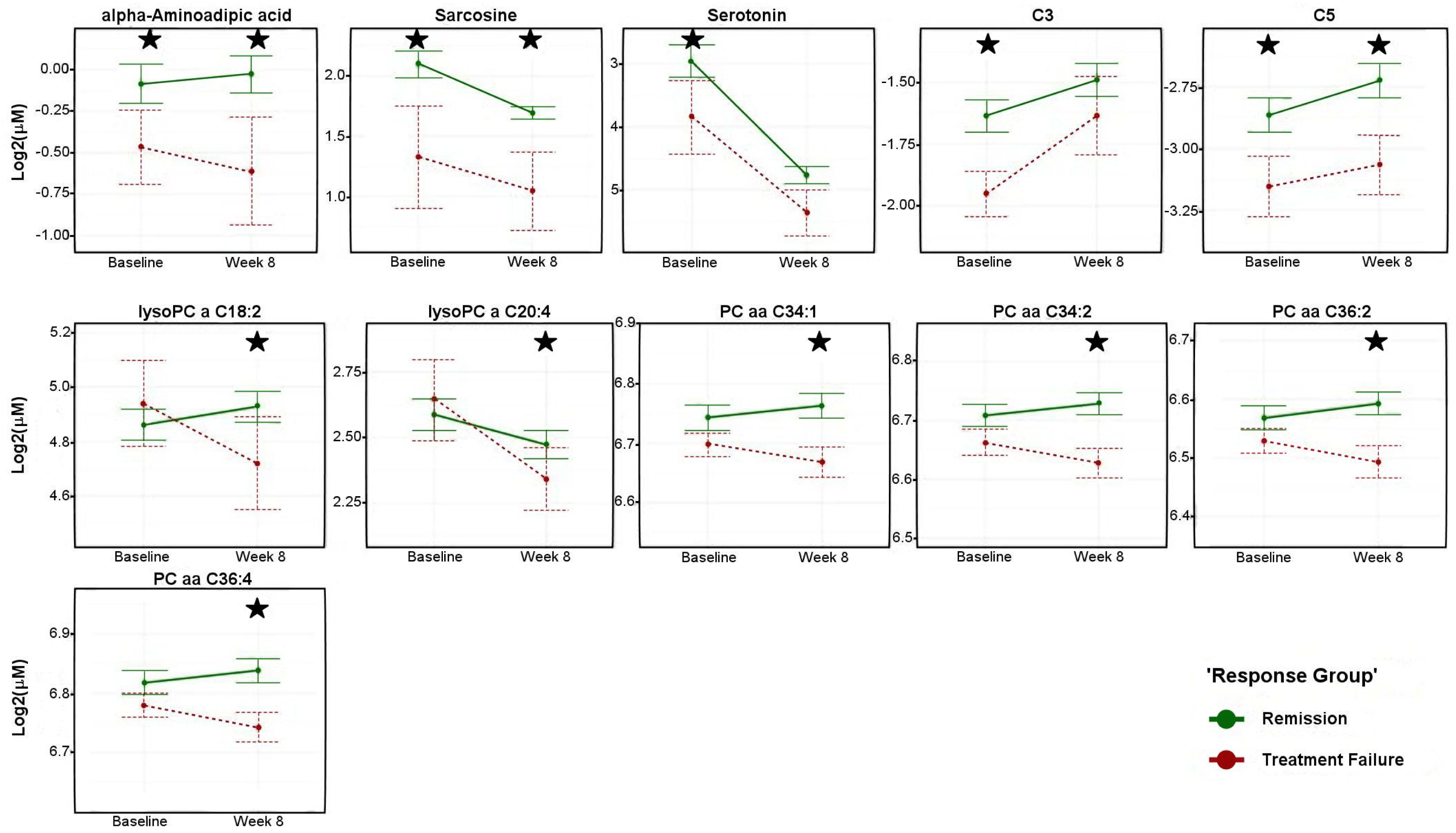
Metabolite Concentrations across Visits by Response Group. Levels (log2 transformed) of metabolites at pre-treatment and 8 weeks post-treatment. Green lines indicate participants who remitted and red lines indicate participants who failed to respond to the treatment at the end of therapy. Asterisks indicate statistical significance of mean differences between the two groups (unadjusted *p*<0.05). Error bars represent standard error of the means. Black stars represents statistical significance at visit. *P*-values were obtained from linear mixed effect models corrected for age, sex, antidepressant and 17-item Hamilton Rating Scale for Depression scores. *Abbreviations*: PC: Phosphatidylcholines, LysoPC: Lyso-Phosphatidylcholines. Metabolite abbreviations are spelled out in Supplementary Table 1.

## Discussion

In the current study, we examined the metabolic consequences of exposure to citalopram/escitalopram in depressed outpatients, thus complementing our previous studies that demonstrated metabolic perturbations in several of the neurotransmitter-related pathways ^68^. Here, we expanded on our previous efforts by focusing on the impact of the SSRI on several classes of metabolites including acylcarnitines, amino acids, biogenic amines, and lipids. In the temporal analyses that assessed changes in metabolite levels from baseline to week 8 in response to the SSRI treatment, significant increases in short-chain acylcarnitines and decreases in the levels of medium- and long-chain acylcarnitines were noted. The levels of several amino acids – such as arginine, proline, tyrosine, citrulline and phenylalanine – were either significantly increased (*q*-value <0.05) or trended toward increase (unadjusted *p*-value <0.05). The biogenic amines like serotonin and sarcosine were significantly decreased while methionine-sulfoxide level increased significantly post-treatment. Among the lipids, several phosphatidylcholines and ether-phosphatidylcholines were increased post-treatment. Moreover, changes in the levels of several lipids and amines were correlated with changes in depression severity (HRSD_17_) after 8 weeks of SSRI treatment. Unique metabolic patterns that differentiate between remitters and participants in the treatment failure group were also noted at baseline and post-treatment. Especially notable were the ether phospholipids that mostly showed an upward trajectory in the remitters while the treatment failure group showed mostly downward trajectories from baseline to week 8 in response to the drug exposure (**Supplementary Fig. 1**).

### Acylcarnitines

Acylcarnitines have been implicated in mitochondrial dysfunction and regulation of energy homeostasis in multiple disease models (Jones, McDonald et al. 2010). They play an important role in brain energy homeostasis and cell signaling cascades (Jones, McDonald et al. 2010) and are used in the diagnosis of several inborn errors of metabolism (Kler, Jackson et al. 1991, Vreken, van Lint et al. 1999, Wanders, Vreken et al. 1999, Cavedon, Bourdoux et al. 2005). Patients with known mitochondrial disorders have frequently reported depressive symptoms (Kato 2001, Fattal, Link et al. 2007, Manji, Kato et al. 2012). Additionally, altered mitochondrial function has been found in patients with a lifetime diagnosis of MDD (Suomalainen, Majander et al. 1992, Gardner, Johansson et al. 2003, Beasley, Pennington et al. 2006).

In our study, after 8 weeks of SSRI treatment, the short-chain acylcarnitines – specifically propionyl carnitine (C3), butyryl/isobutyryl carnitine (C4) and isovaleryl/methylbutyryl carnitine (C5) – were significantly increased while acetyl carnitine (C2) levels were decreased. Several studies have previously reported perturbations in C2 after antidepressant treatment in depressed patients ^20, 32, 52^. In a rat model of depression, incomplete β-oxidation of fatty acids has been associated with elevated medium- and long-chain acylcarnitines (Chen, Wei et al. 2014). This finding is in line with our observation of decreased medium- and long-chain acylcarnitines after 8 weeks of anti-depressant therapy, and suggests that the drug may act to restore the mitochondrial β-oxidation process with greater utilization of the medium- and long-chain acylcarnitines. Remarkably, in our study, the plasma ratio of long-chain acylcarnitines to free carnitine (C16:0 +C18:0/C0) (which is a marker for the activity of carnitine palmitoyltransferase 1, the rate-limiting step in the uptake of fatty acids into mitochondria ^69^) was significantly reduced at week 8 (*p*-value < 3.4E-06). In two recent studies that involved schizophrenia patients, similar patterns of short-, medium- and long-chain acylcarnitine perturbations were observed ^34, 35^ and antipsychotic treatment significantly reduced levels of C2, increased the short-chain acylcarnitines, and decreased the levels of medium- and long-chain acylcarnitines ^35^. Taken together, these findings suggest a similar pattern of mitochondrial dysfunction and a drug-induced functional restoration thereof across the spectrum of neuropsychiatric disorders.

We also found that the changes in short-chain acylcarnitines (C3, C4, C5) over 8 weeks of SSRI treatment were correlated with changes in branched-chain amino acids (BCAAs; isoleucine and valine). C3 and C5 acylcarnitines are products of the degradation of BCAAs ^70^. We and others ^21, 71^ have previously shown that perturbations in plasma BCAA levels are significantly associated with MDD. BCAAs have well-established anabolic effects on protein metabolism that involve activation of the mTOR pathway ^21^. Baranyi et al. ^71^ suggested that the decrease of BCAAs in depressive patients could result in the dysregulation of the mTOR pathway, leading to depressive symptomology and lower energy metabolism. In our study, we also observed that BCAA levels increased with the antidepressant exposure, though not significantly, and that remitters had higher baseline BCAA levels that further increased post-treatment compared to the treatment failure group.

### Amino Acids and Biogenic Amines

After 8 weeks of SSRI treatment, we noted significant perturbations in the urea cycle and nitric oxide cycle metabolites. Besides clearance of waste nitrogen, the distinct biochemical goals of these cycles involve production of the intermediates ornithine, citrulline, and arginine for the urea cycle; and polyamine production and the production of nitric oxide for the nitric oxide pathway ^72^ (**Fig. 1B**). In our study, after 8 weeks of drug exposure, plasma arginine levels increased significantly, while ornithine levels showed a trend to be lower compared to baseline. The ratios of citrulline/arginine (*p*-value < 0.013) and asymmetric dimethyl arginine/arginine (ADMA/Arg, *p*-value < 8.62E-05) were significantly lower compared to baseline, both of which may indicate potential increases in activity of Nitric Oxide Synthase and increased production of Nitric Oxide (NO) post-treatment. NO is a known modulator of the major neurotransmitters – norepinephrine, serotonin, dopamine and glutamate – involved in the neurobiology of MDD ^73^. Rotroff and colleagues have found a negative association between the levels of ornithine and citrulline with changes in the Montgomery-Åsberg depression rating scale after treatment with ketamine ^20^. In line with our findings, several studies have reported lower levels of citrulline in MDD patients compared to healthy controls ^51, 52^, while another reported lower levels of arginine in MDD patients compared to healthy controls ^53^. Hence, a full analysis of these pathways is warranted in which measurements of more metabolites are covered.

Similar to our previous findings ^68^, the biogenic amine serotonin decreased significantly with the drug exposure. A recent study from our group found that after 4 weeks of SSRI exposure, the decrease in sarcosine levels was correlated with the decrease in serotonin levels ^21^. Sarcosine is an endogenous amino acid involved in one-carbon metabolism and has recently emerged as a promising therapy for schizophrenia, acting as an N-methyl-D-aspartate receptor agonist ^74^. Sarcosine and serotonin were also highly correlated to each other in our study. Additionally, at baseline, their levels were significantly higher in the remitters compared to the treatment failure group and remained higher even at the end of treatment. This suggests that further studies are warranted to determine whether their higher baseline levels were critical to the drug response.

Among the other members of this class, changes in histidine and kynurenine levels from baseline to week 8 were inversely associated with changes in depressive symptoms scores (HRSD_17_). Both of these metabolites have been previously implicated in the pathophysiology of depression ^45, 75^. We also found significant increases in methionine sulfoxide (MetSO; **Fig. 2A**) and the ratio of methione sulfoxide/methionine (MetSO/Met; *p*-value < 0.0008) from baseline to week 8. MetSO is a primary oxidation product of methione, and is a possible biomarker for oxidative stress ^46^ which has already been associated with MDD. Interestingly, a few studies have suggested that methione residues on proteins can act as sacrificial antioxidants, and that the reversible conversion of methione to MetSO may act as a reversible redox switch to regulate the function of proteins ^76–78^.

### Lipids

Among the lipids studied, the perturbations amongst the phosphatidylcholines (PCs) containing either the diacyl or the alkyl-acyl moieties were strong, and many of them were inversely associated with changes in the depressive symptom scores from baseline to week 8 (**Fig. 2A**, **Supplementary Table 2**). Furthermore, upon stratification between remitters and participants with treatment failure, several interesting trends emerged. First, the lysophosphatidylcholines containing a single fatty acyl chain (lysoPCs -C16:0, -C17:0, -C18:0, - C18:1, C18:2, C20:3 and C20:4) mostly showed downward trajectories from baseline to week 8 among the treatment failure group but not so among the remitters. Second, several diacyl PCs that showed inverse correlations to changes in depression scores (**Fig. 3 and Supplementary Table 2**) (e.g., PC aa –C34:1, 34:2, 36:2, 36:3, 36:4, 38:4) showed very similar trajectories in that the remitters all had higher baseline levels that increased further at week 8, compared to the treatment failure group which showed very similar downward trajectories (**Fig. 3**). Lastly, the ether-phosphatidylcholines as a group showed a distinct pattern of perturbation: the treatment failure group mostly had significantly higher baseline levels compared to the remitters, and these metabolites then followed downward trajectories over the treatment period whereas the remitters either showed an increase or stayed unchanged (**Supplementary Fig. 1**). Overall, the lipids displayed remarkable differences between the remitters and treatment failure group regarding how the two groups responded to the drug. While it is difficult to come up with plausible explanations for such observed differences, there seems to be an important role for these lipids in contributing to drug response.

Lipids are involved in crucial brain functions, including cell membrane structure, membrane transmitters, energy metabolism, and neuroendocrine function ^54^. Blood lipid profile changes have been implicated with the pathogenesis of depression, schizophrenia, and Alzheimer’s disease ^55–59^. The association between the changes in lipid profiles, depression and anxiety has been previously reported in several studies ^79–83^.

Of special interest is the distinct pattern we observed among the ether phospholipids. The distinctive chemical feature of the ether lipids is the ether bond at the *sn*-1 position of the glycerol backbone where a fatty alcohol is attached as opposed to the more common diacyl moiety containing phospholipids. The emerging role of these lipids in neurological diseases like Alzheimer’s Disease and autism is currently garnering much interest ^84^. They are essential in shaping membrane integrity and properties, thereby potentially impacting numerous biological processes and functions, serve as antioxidants, and have recently been shown to be critical modulators of neurotransmitter homeostasis and release as well as behavioral abnormalities in a knock-out mouse model ^85 86^. The authors reported reduced brain levels of various neurotransmitters in the ether-lipid-deficient knock-out mice, which they attributed to altered synaptic vesicles releasing reduced amounts of neurotransmitters. Recently, Knowles et al. demonstrated a shared genetic etiology between MDD and ether-phosphatidylcholine species that contain arachidonic acid, the latter being a precursor to the pro-inflammatory prostaglandins^87^. The initial stages of ether-lipid synthesis occur in the peroxisomes, including the rate-limiting enzymatic processes. It is also possible that peroxisomal disorders and subsequent metabolic resilience result in the distinct patterns observed in the trajectories of ether-phospholipids in remitters vs. those in the treatment failure group.

To summarize, our data suggest that SSRI treatment results in changes in the acylcarnitine, lipid and amino acid profiles. The changes in acylcarnitine profiles were similar to a handful of studies conducted in other psychiatric illnesses, such as first-episode psychosis, which suggests a substantial overlap of implicated pathways and mechanisms across mental illnesses. The data show that after treatment, long-chain acylcarnitines are reduced but short-chain acylcarnitines are increased. The reduced medium- and long-chain fatty acylcarnitine levels could have resulted from enhanced mitochondrial fatty acid oxidation after treatment. They could also have resulted from an inhibition of fatty acid transport to mitochondria. Increased short-chain acylcarnitines – including C3 and C5 – support the latter case since the majority of these short-chain acylcarnitines are produced from amino acids, indicating an energy switch from fatty acid oxidation to other energy substrates. The reduced content of acetylcarnitine suggests the existence of a possible starvation state of energy, which further supports the latter case. However, further studies are needed to definitively clarify these possibilities. Whatever the case, it appears that reduction of ATP production is likely the consequence of the drug treatment. This finding could be further explored to develop new therapeutics targeting other approaches to reduce ATP availability. More importantly, a reduced CPT1 activity could lead to the accumulation of palmitate, oleate, and their relevant fatty acylCoA species since CPT1 is relatively selective for transporting these two fatty acyls to mitochondria.

We also noted perturbation of phospholipid profiles. Additionally, remitters show unique changes in their lipid profiles compared to participants with treatment failure. Although only countable PC species are significantly changed in the remitter group, essentially all PC species containing saturated or monounsaturated fatty acyl chains (i.e., PC species containing ≤3 total double bonds) tended to increase, whereas PC species containing ≥4 total double bonds were reduced or tended toward reduction. These changes were not observed in the treatment failure group. These types of changes likely resulted from the remodeling of PC species, likely in the liver, through a combined action of phospholipases (e.g., phospholipase A2) and acyltransferase activities. Although the activity of lecithin cholesteryl ester transferase could also contribute to this change, this activity yields less selective remodeling of fatty acyl chains.

The changes in the acylcarntines and PC species in the remitter group could be inter-related. For example, the accumulation of palmitate, oleate, and their relevant fatty acyls due to a reduced CPT1 activity as discussed above could lead to maladaptive changes of the PC pattern due to the precursor availability for an acyltransferase activity as uncovered in this study. Alternatively, remodeling of PC species due to the action of SSRI treatment may produce changes of membrane structure and function, thus affecting the activities of membrane proteins such as CPT1. However, the cause-and-effect relationship remains unknown. Further studies to clarify this relationship – such as measuring the inhibitory effect of SSRI on CPT1 activities and determining the accumulation of specific acylCoAs – are clearly needed.

The above information shows that assessing metabolic profiles may enable the mapping of global biochemical changes in MDD, provide a means to characterize the remitted and depressed states, and provide a way of characterizing the effects of antidepressants on metabolic pathways. This approach may ultimately inform therapeutic choices, thus reducing trial-and-error prescribing and contributing to personalizing therapeutic treatment for patients with MDD.

This study has several limitations. First, some of the investigated acylcarnitines – especially dicarboxylic and hydroxylated species (C3-DC, C5-DC, C5-MD-C, C16:OH, C18:OH) – are recognizably high only in patients with rare inborn errors of metabolism (**Supplemental Table 1**), and display undetectable concentrations in most individuals. In this report, the flow-injection MS/MS method used to confirm exact structures for low-level species may lack molecular specificity. Nevertheless, the measures reported here show exceptional technical reproducibility. Of interest, although we investigated many of the low-abundance (i.e. <0.1 µM) acylcarnitines, our average observed values for each analyte in the cohort are below the pathological clinical reference threshold described by Mayo Clinic Laboratories ^88^. Moreover, the low-abundance acylcarnitines reported here may benefit from utilization of an assay with greater molecular specificity to confirm the exact molecular speciation, such as a research-grade LC-MS/MS assay reported previously ^89^. Another limitation is the relatively small sample size. It will be necessary to replicate and validate our findings in larger independent studies. Further, the lack of a control group limits our ability to control for spontaneous changes in the acylcarnitine profiles that occur regardless of treatment or the passage of time. Diet, medication and other factors may also contribute to changes in metabolite levels over time. Finally, due to small sample size, our analyses were not stratified by sex or other confounding factors.

In conclusion, metabolomics provides powerful tools for understanding disease mechanisms, the molecular basis of disease heterogeneity, and variation of response to treatment. The altered metabolic profiles suggest that mitochondrial energetics – including acylcarnitine metabolism, transport and its link to β-oxidation, as well as lipid membrane remodeling – are implicated in recovery from a depressed state. If validated in large studies, these approaches could provide molecular signatures that help explain the variability in response to antidepressant treatment.

## Acknowledgements

The authors are grateful for the support of NIH, to Lisa Howerton for her administrative support, to David S. Millington, PhD for insightful discussions and suggestions and to the study participants and their families of the Mayo Pharmacogenomics Research Network-Antidepressant Pharmacogenomics Medication Study (PGRN-AMPS). The research and the authors are supported by funding from the NIH. This work was funded by grant support to Rima Kaddurah-Daouk through NIH grants R01MH108348, R01AG046171 & U01AG061359, RF1AG051550. R.M.W. was supported by NIH grants RO1 GM28157, U19 GM61388, U54 GM114838, and NSF1624615. SB was supported by the NIH grant R01MH108348.

## SUPPLEMENTARY MATERIALS

**Supplementary Fig 1.**
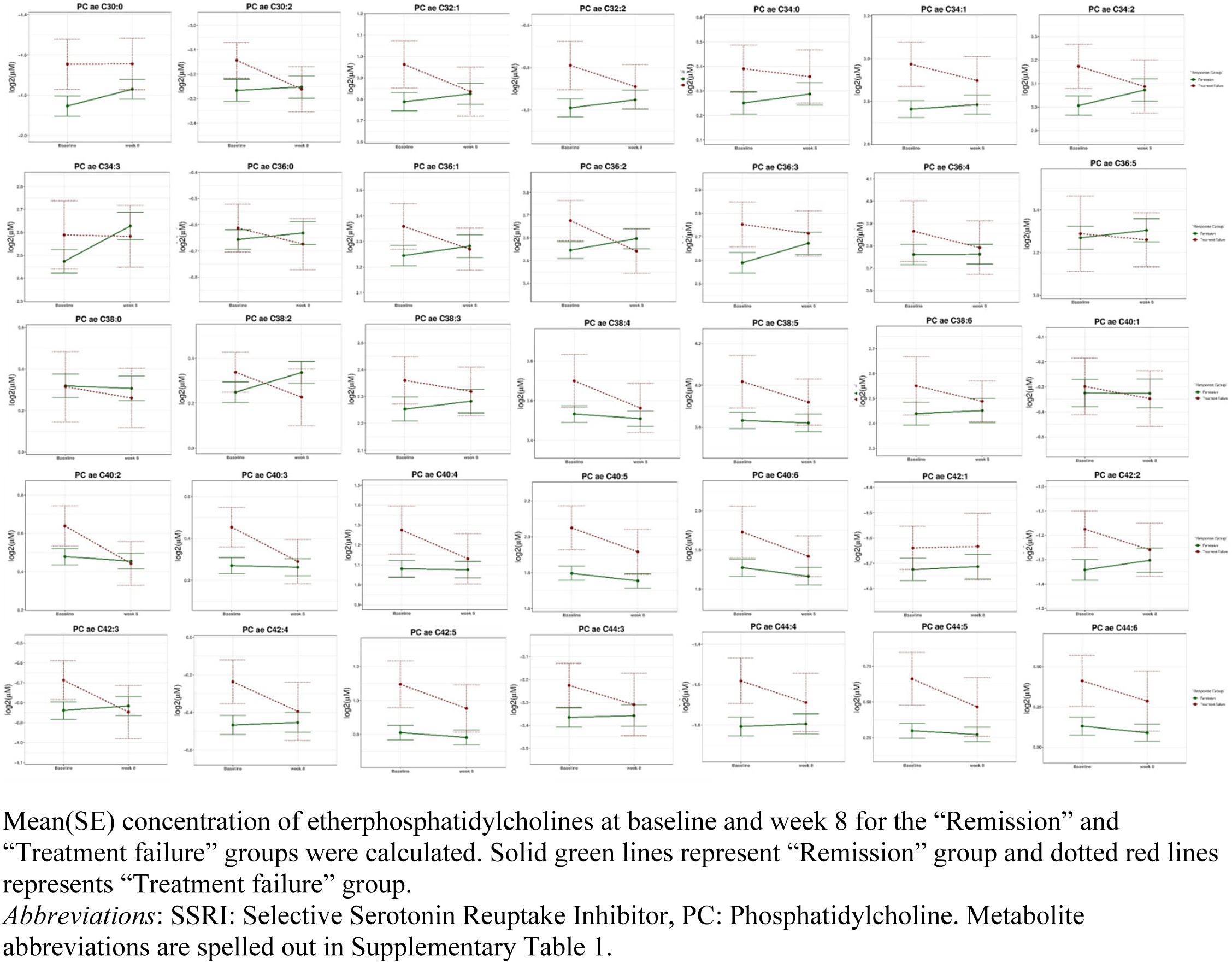
Effect of 8 Weeks SSRI Treatment on Levels of Ether Phosphatidylcholines.

**Supplementary Table 1.**
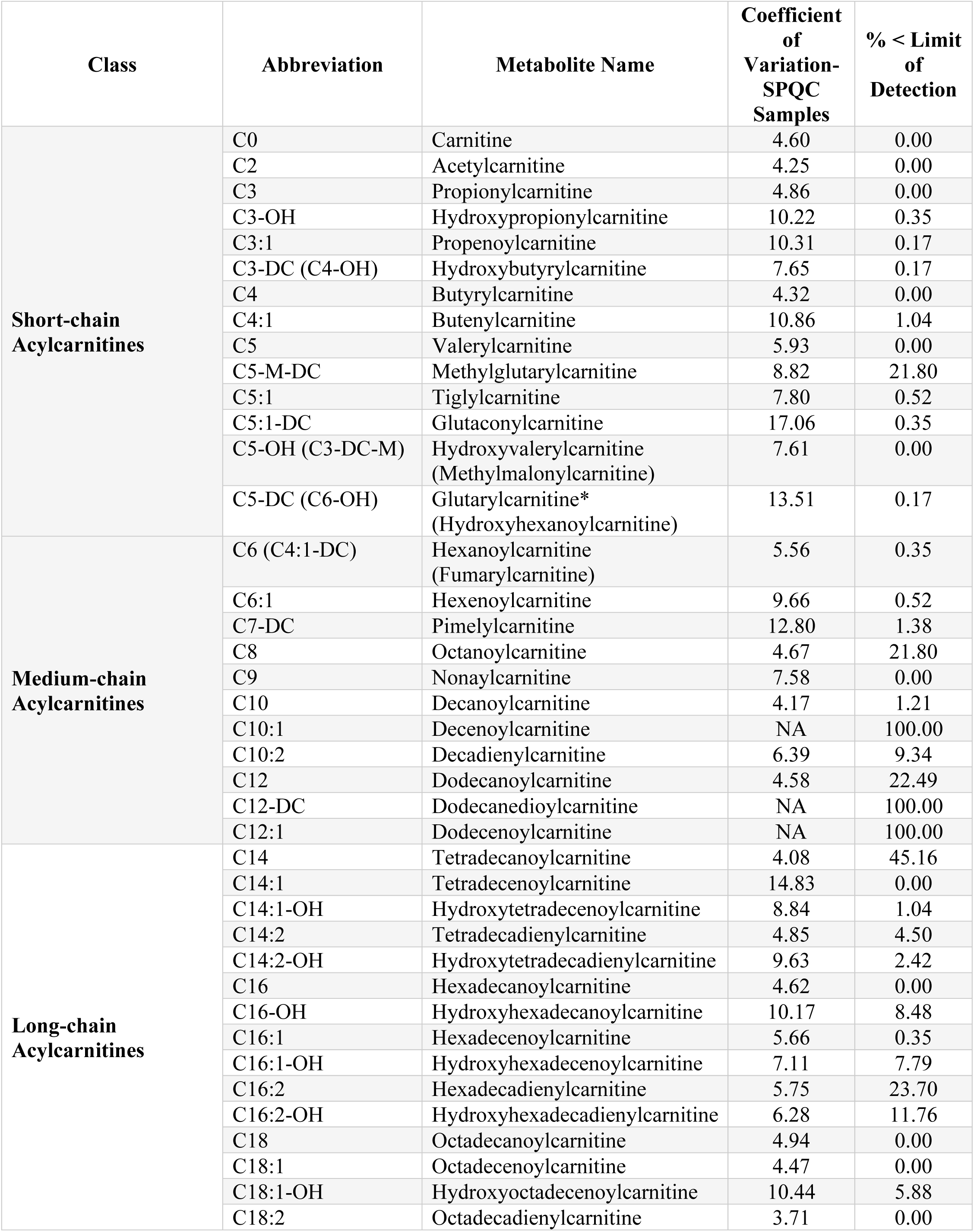
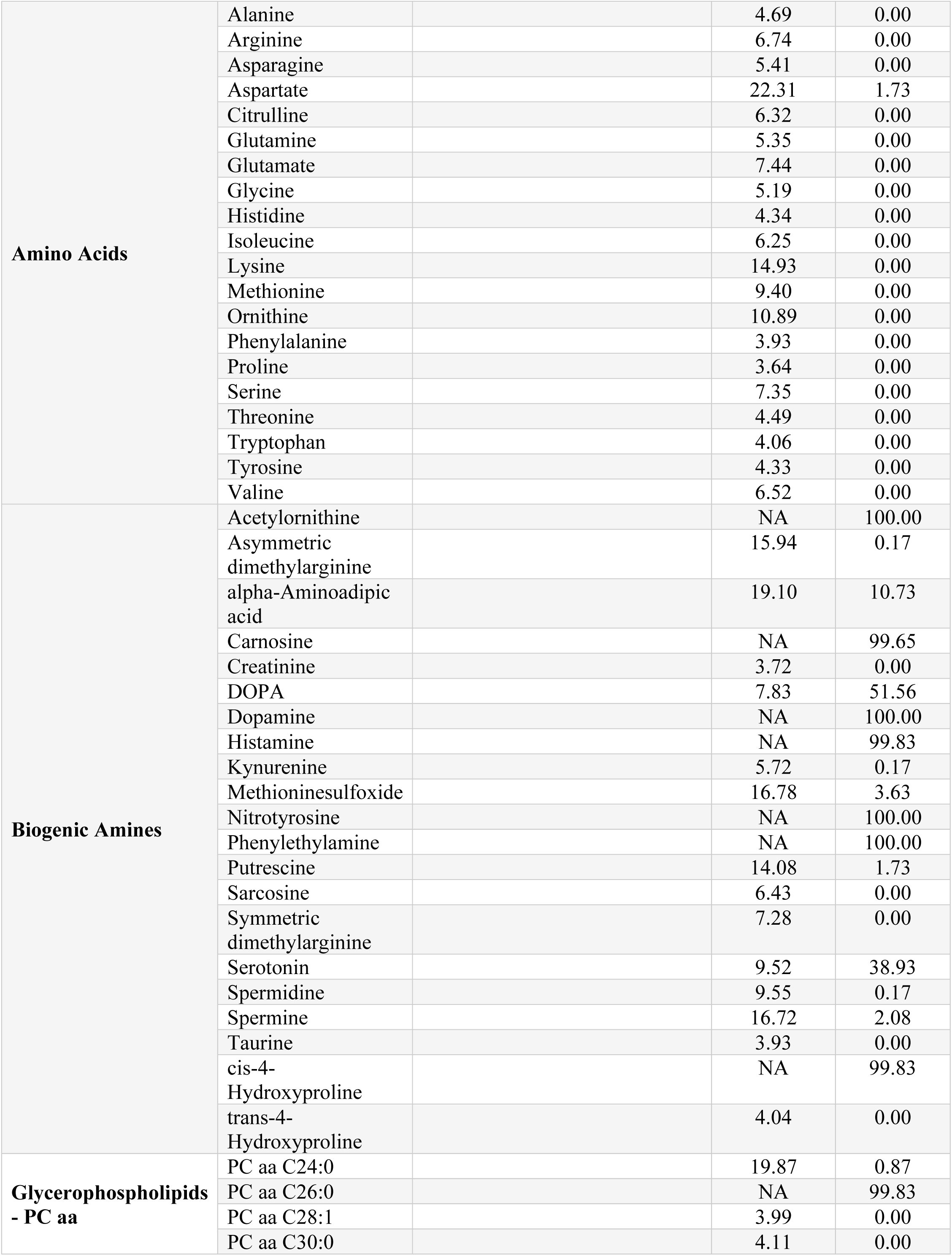
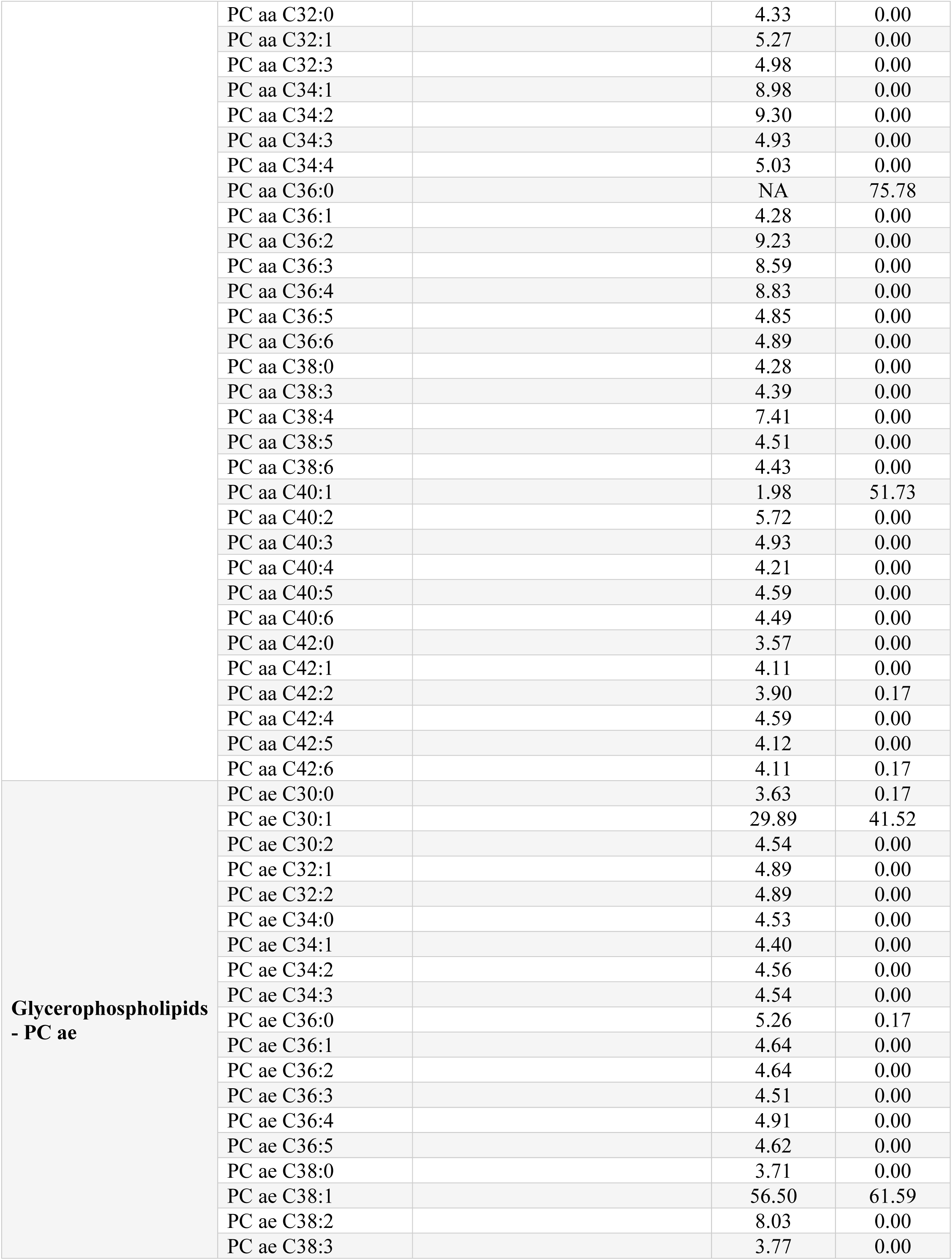
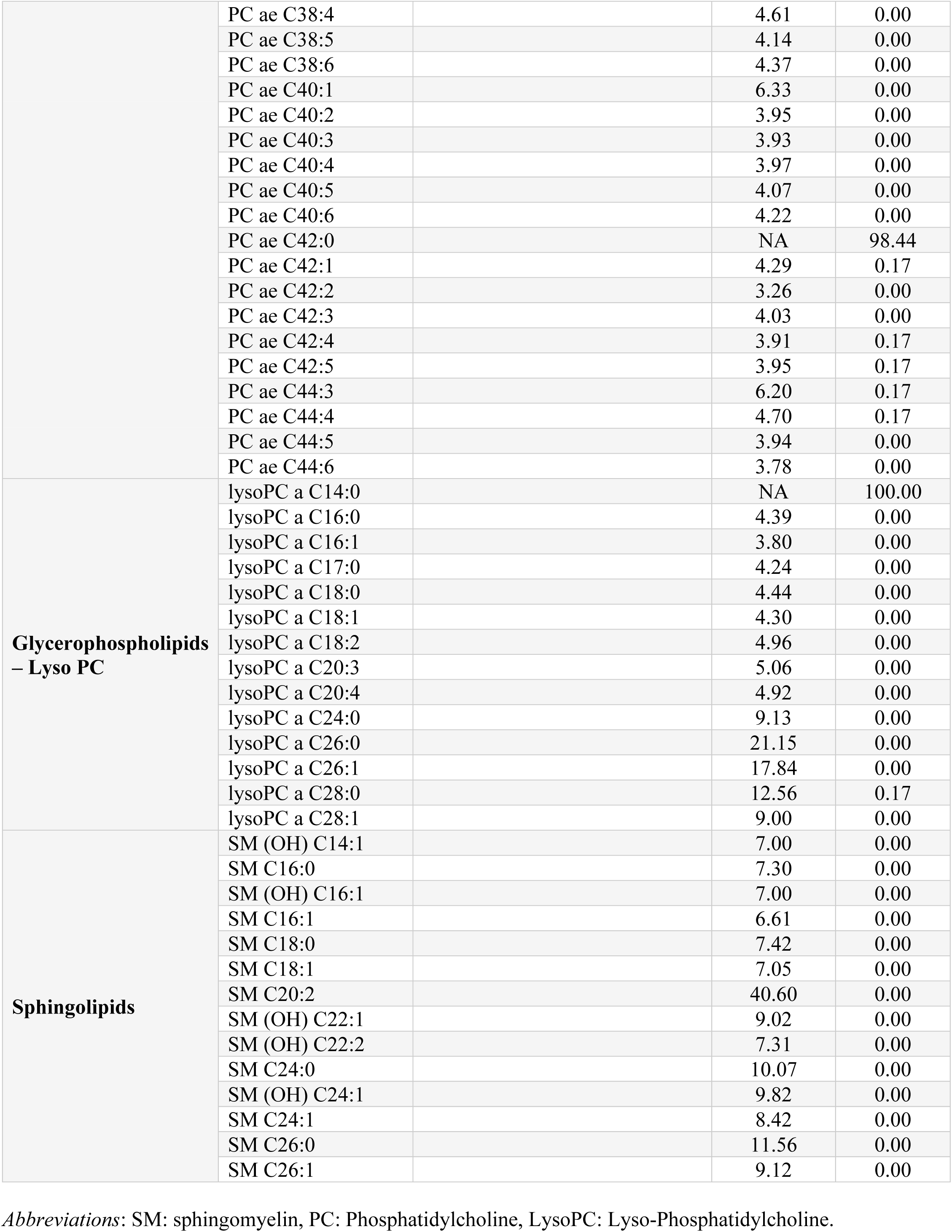
Metabolites Measured by Biocrates P180^®^ Kit

**Supplementary Table 2.**
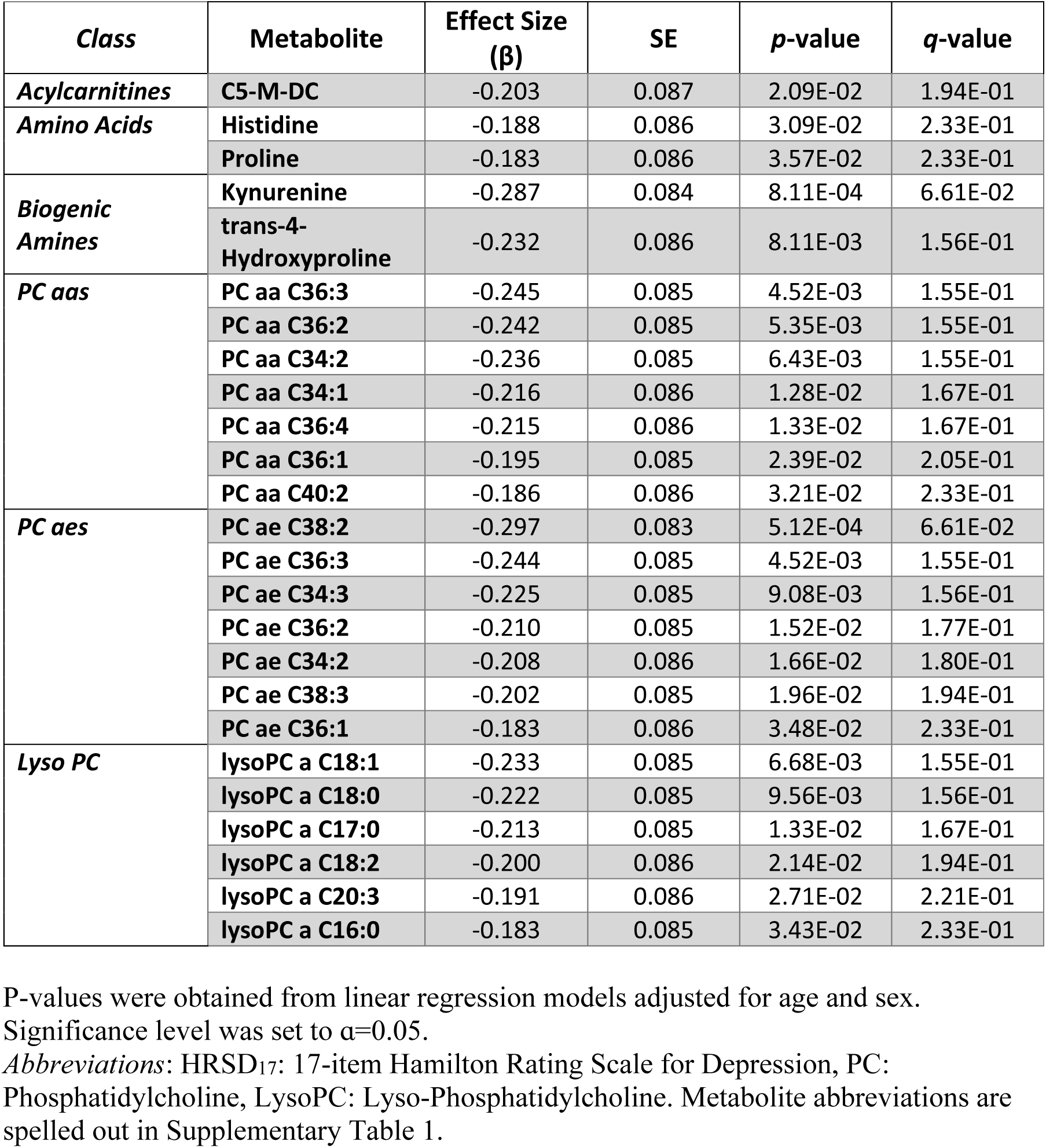
Significant Associations Between Change in HRSD_17_ Score and Change in Metabolite Levels Over 8 Weeks of Treatment

**Supplementary Table 3.** Differences in Metabolites Levels between Remitters and Treatment Failures across Visits

| Baseline |  |  | 8 weeks |  |  |
| --- | --- | --- | --- | --- | --- |
| Metabolite | Estimated Log<sub>2</sub> Difference (SE) | *p*-value | Metabolite | Estimated Log<sub>2</sub> Difference (SE) | *p*-value |
| alpha-Aminoadipic acid | 0.54 (0.26) | 3.96E-02 | alpha-Aminoadipic acid | 0.58(0.26) | 2.68E-02 |
| Sarcosine | 0.57(0.25) | 2.43E-02 | Sarcosine | 0.52(0.25) | 3.85E-02 |
| Serotonin | 1.02(0.48) | 3.24E-02 | C5 | 0.38(0.15) | 1.24E-02 |
| C3 | 0.35(0.15) | 2.35E-02 | lysoPC a C18:2 | 0.34(0.16) | 3.60E-02 |
| C5 | 0.31(0.16) | 3.91E-02 | lysoPC a C20:4 | 0.28(0.14) | 4.25E-02 |
|  |  |  | PC aa C34:1 | 0.09(0.04) | 4.00E-02 |
|  |  |  | PC aa C34:2 | 0.10(0.04) | 1.55E-02 |
|  |  |  | PC aa C36:2 | 0.10(0.04) | 1.88E-02 |
|  |  |  | PC aa C36:4 | 0.10(0.04) | 2.94E-02 |

Linear mixed effect models were used to assess the significance of visit by treatment outcome interaction. Models were adjusted for age, sex, and baseline HRSD<sub>17</sub> score. Significance level was set to $\alpha=0.05$ .
*Abbreviations:* LysoPC: Lyso-Phosphatidylcholine. PC: Phosphatidylcholine. Metabolite abbreviations are spelled out in Supplementary Table 1.

